# Heat Shock Proteins Function as Signaling Molecules to Mediate Neuron-Glia Communication During Aging

**DOI:** 10.1101/2024.01.18.576052

**Authors:** Jieyu Wu, Olivia Jiaming Yang, Erik J. Soderblom, Dong Yan

## Abstract

The nervous system is primarily composed of neurons and glia, and the communication between them plays profound roles in regulating the development and function of the brain. Neuron-glia signal transduction is known to be mediated by secreted or juxtacrine signals through ligand-receptor interactions on the cell membrane. Here, we report a novel mechanism for neuron-glia signal transduction, wherein neurons transmit proteins to glia through extracellular vesicles, activating glial signaling pathways. We find that in the amphid sensory organ of *Caenorhabditis elegans*, different sensory neurons exhibit varying aging rates. This discrepancy in aging is governed by the crosstalk between neurons and glia. We demonstrate that early-aged neurons can transmit heat shock proteins (HSP) to glia via extracellular vesicles. These neuronal HSPs activate the IRE1-XBP1 pathway, further increasing their expression in glia, forming a positive feedback loop. Ultimately, the activation of the IRE1-XBP-1 pathway leads to the transcriptional regulation of chondroitin synthases to protect glia-embedded neurons from aging-associated functional decline. Therefore, our studies unveil a novel mechanism for neuron-glia communication in the nervous system and provide new insights into our understanding of brain aging.

## Introduction

Aging causes a decline in brain function and is the most significant risk factor for neurodegenerative diseases^1–3^. Over the past few decades, extensive efforts have been dedicated to studying the aging of neurons, leading to a significant understanding of molecular mechanisms underlying neuronal aging^4^. However, it is worth noting that more than half of the cells in the brain are glia^5^, and the function significance of glia and their crosstalk with neurons during aging are not well understood.

With increasing evidences showing that glial cells are versatile regulators in various aspects of brain function previously attributed solely to neurons, substantial knowledge of neuron-glia interactions has been unveiled in the past three decades^6^. The crosstalk between neurons and glia presents great complexity and heterogeneity throughout the nervous system. Glial cells can sense different aspects of neuronal activity, such as electrical signals, synaptic neurotransmitters, structural changes, and energy demands, through a cassette of membrane receptors, channels, transporters, and pumps^7–9^. In turn, glial cells can release different molecular signals, such as gliotransmitters (glutamate, ATP, GABA and D-serine) and cytokine (TNFα, IL-10 and TGFβ), to impact neuronal activity^7,10,11^. Thus, neuron and glia can sense each other to respond different stimulations through cell-cell signal transduction, thereby regulating brain function. Besides functional regulation, the interactions between neurons and glia also shape their development^12–14^. In all these instances, the signal transductions between neurons and glia rely on ligand-receptor interactions on the cell membrane. Yet, it remains to be explored whether neurons and glia can communicate through other mechanisms in physiological and pathological conditions.

Extracellular vesicles (EVs) are membrane-surrounded structures released by nearly all cells. EVs exhibit heterogeneity and contain various contents including proteins, lipids, nucleic acids, metabolites, and even organelles^15,16^. The contents of EVs are believed to be selectively packaged rather than passively included, leading to the proposal that EVs play a role in mediating cell-cell communications. Extensive research has elucidated the functions of EVs in cell-cell communications, particularly in virus infections such as Epstein-Barr virus (EBV) infections. Studies demonstrate that virus-infected cells can pack viral miRNA, mRNA, and lncRNA into EVs, transmitting them into healthy cells and causing further damages^17,18^. Additionally, EVs are implicated in cancer progression via transfer of EV-associated miRNAs, which is sufficient to driving the transformation of non-tumorigenic cells into a tumor^19–21^. In the nervous system, EVs are found to be released from all major cell types, including neurons, astrocytes, microglia, and oligodendrocytes^22–24^. However, our understanding of the function of EVs in the brain is still limited. Notably, one of the most characterized neuronal regulations involving EVs comes from the study of a protein called Arc. Arc can form virus-like capsids to pack its mRNA into EVs to mediate neuron-neuron transmission, and the transmitted mRNA can influence the function of neurons^25^. Other studies indicate that neuronal and glial EVs can impact neuronal survival and synaptic activity in vitro, though the mechanisms through which EVs mediate these processes remain unclear^26–28^. To date, most studies of EVs in cell-cell signal transduction have focused on RNA as signal components. However, beyond RNA, many proteins are packaged into EVs, and the functions of these proteins in signal transduction, specifically in the brain, remain to be studied.

In this study, we demonstrate that early aged neurons can transmit heat shock proteins (HSP-4) into glia through EVs. These neuronal HSP-4 proteins activate the glial IRE1-XBP1 pathway, further increasing HSP-4 expression in glia and forming a positive feedback loop. Consequently, the activation of the IRE1-XBP1 pathway promotes the transcriptional regulation of chondroitin synthases to protect glia-embedded neurons from aging-associated functional decline. Our results uncover a novel mechanism for neuron-glia communication during aging, wherein neurons directly transmit proteins into glia to regulate glial function, thereby protecting other neurons.

## Results

### Varied aging rates among sensory neurons are intrinsically correlated

To investigate neuron-glia interactions during aging, we used the *C. elegans* amphid sensory organ as a model^29^. The amphid sensory organ in *C. elegans* comprises 12 sensory neurons and 2 glial cells (socket glia and amphid sheath (AMsh) glia)^30^. It exhibits structure and morphology similar to sensory units in other species, such as mammalian taste buds and olfactory bulbs^31^. These 12 neurons can be further classified into AMsh- channel neurons and AMsh glia-embedded neurons, and their dendritic receptive endings are wrapped in a channel formed by AMsh glia to be exposed to the external environment and are entirely embedded within AMsh glia in a hand-in-glove configuration, respectively^30^ (Figure 1A). AMsh glia are essential for the function of the sensory organ and share developmental and functional similarities with mammalian glia^29,32^. To study how aging affects the function of the amphid sensory organ, we began by testing the function of two AMsh-channel neurons, ASH and ADL neurons, and two AMsh glia-embedded neurons, AWA and AWC neurons, during aging (Figure 1A). ASH/ADL neurons mediate repulsion behaviors in response to octanol, while AWA and AWC neurons sense 2-methylpyrazine and benzaldehyde, respectively, to initiate attraction behaviors^32,33^ (Figure S1A and S1B).

**Figure 1.**
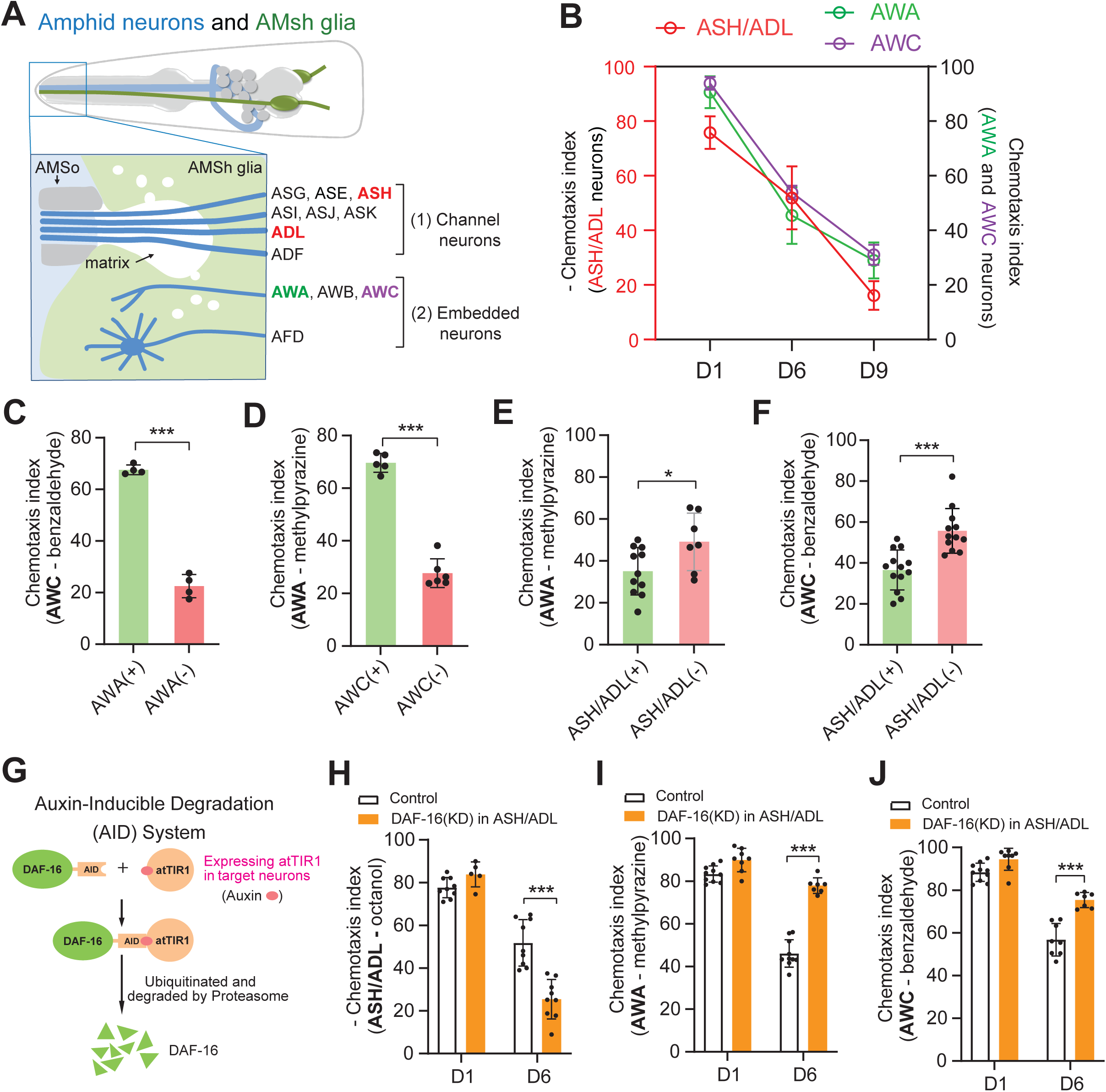
AMsh-channel and AMsh glia-embedded neurons age differently in individual animals. **(A)** Schematic diagram of the amphid sensory organ of *C. elegans*. (1) AMsh-channel neurons; (2) AMsh glia-embedded neurons. **(B)** Results from chemotaxis experiments show that ASH/ADL, AWA and AWC neurons exhibit age-dependent functional decline in response to octanol, 1% 2-methylpyrazine and 0.5% benzaldehyde, respectively. D1, D6 and D9 represent the adult stage day 1, day 6, and day 9 animals. Data are shown as mean ± SD **(C** to **F)** Function correlation analyses among ASH/ADL, AWA and AWC neurons. After the initial chemotaxis assay testing, day 6 animals were divided into two groups (Figures S1A and S1B): animals that respond to corresponding odorants were labeled as (+); and animals that did not respond to corresponding odorants were labeled as (−). The results from chemotaxis assays for different sensory neurons by using these two groups of animals were presented in **C** to **F**. Date are shown as mean ± SD. Student’s *t*-test, *P < 0.05, ***P < 0.001. **(G)** Schematic of Auxin-inducible DAF-16 degradation in target neurons ^43^. To exclusively deplete DAF-16 in ASH/ADL neurons, we generated a transgene expressing atTIR1 (derive from Arabidopsis thaliana) under an ASH/ADL-specific promoter, P*gpa-11*, in *daf-16(ot853)* animals. *daf-16(ot853) [daf-16::linker::mNeonGreen::3xFlag::AID]* is a knockin strain in which a degron-tagged mNeonGreen is inserted at the C-terminus of *daf-16*. *daf-16(ot853)* without atTIR1 expression was used as a control. Adult animals grow on the NGM plates supplied with 5 μM auxin (indole-3 acetic acid, sigma). All DAF-16 depletion experiments were carried out using this setting. **(H** to **J)** Chemotaxis assays for ASH/ADL (**H**), AWA (**I**) and AWC (**J**) neurons using octanol, 1% 2-methylpyrazine, and 0.5% benzaldehyde, respectively, in day 1 (D1) and day 6 (D6) animals with (DAF-16 (KD)) and without (Control) DAF-16 depletion in ASH/ADL neurons. Data are shown as mean ± SD. Student’s *t*-test, ***P < 0.001.

The average lifespan of *C. elegans* is approximately 2 weeks, and most animals exhibit noticeable neuronal aging phenotypes within the first 9 days^34–37^. Therefore, we used the adult stage day 1 (D1), day 6 (D6), and day 9 (D9) animals to assess the function of amphid sensory neurons during aging. Utilizing classic chemotaxis assays^32,38^, we observed age-dependent functional decline in ASH/ADL, AWA, and AWC neurons (Figures 1B, S1A, and S1B). While from a population perspective these neurons appeared to age collectively at a similar rate, this data did not address the relationship among the aging of these sensory neurons in individual animals. To address this question, following the initial chemotaxis testing, we first divided day 6 animals into two groups, with and without responses, and then compared the function of other sensory neurons between these two groups (Figures S1A and S1B). We found that the function of AWA and AWC neurons were tightly correlated with each other, that animals with functional AWA/AWC neurons were more likely to have normal AWC/AWA functions when compared with those without AWA/AWC functions (Figures 1C and 1D). In contrast, the function of ASH/ADL neurons were negatively correlated with AWA/AWC functions, that animals without ASH/ADL functions tended to have functional AWA/AWC neurons when compared to those with ASH/ADL functions (Figures 1E and 1F). However, the functionality of ASH/ADL neurons could not be predicted by AWA or AWC functions, as animals with or without AWA/AWC functions performed similarly when tested for ASH/ADL functions (Figures S1C and S1D). This unilateral negative correlation between ASH/ADL and AWA/AWC functions during aging suggests that the underlying signals may be initiated by ASH/ADL neurons to affect AWA/AWC neurons.

To uncover the mechanisms behind the negative correlation between ASH/ADL and AWA/AWC functions in day 6 animals, we examined whether the aging of ASH/ADL neurons could protect AWA/AWC neurons. In *C. elegans*, one of the major aging signals is the insulin-like growth factor signaling pathway, and its downstream transcription factor *daf-16* can regulate neuronal aging cell autonomously^39,40^. Using a well-established *daf-16* knockin translational reporter^41^, we observed the expression of *daf-16* in ASH/ADL neurons, while it was absent in the nearby AMsh glia (Figures S1E and S1F). To induce ASH/ADL neuronal aging, we employed auxin-inducible degradation system (AID) to deplete DAF-16 proteins exclusively in ASH/ADL neurons^42,43^ (Figure 1G). We found that selectively depletion of DAF-16 in ASH/ADL neurons accelerated their aging and caused further decline in ASH/ADL functions when compared to the same age control animals (Figure 1H), and these modulations largely prevented the functional decline of AWA/AWC neurons caused by aging (Figures 1I and 1J). Taken together, we believe that the aging of ASH/ADL neurons can prevent aging-associated functional decline of AWA/AWC neurons.

### Neuronal aging activates the UPR^ER^ pathway in AMsh glia

Although ASH/ADL and AWA/AWC neurons do not directly communicate with each other, they both make contact with the same glial cell, namely AMsh glia (Figure 1A). Since bidirectional regulations between neurons and glia have been well-documented in many organisms^8,10^, we investigated whether AMsh glia are involved in the regulation between ASH/ADL and AWA/AWC neurons. First, we examined how aging affected AMsh glia and found that AMsh glia in aged animals displayed aging-associated features, such as accumulations of large vacuole structures (Figures S2A and S2B). Further examinations of these vacuole structures showed that they were co-localized with the late-endosome marker RAB-7 and the lysosome marker LMP-1^44,45^ (Figure S2C), similar to what has been observed in aged neurons^46–48^. Using these abnormal vacuole structures as a marker, we evaluated the relationship between AMsh glial aging and the function of sensory neurons. We observed a positive correlation between the aging of AMsh glia and the aging-associated functional decline of AWA/AWC neurons, while a negative correlation was noted with ASH/ADL functional decline (Figures S2D-S2F). These findings suggest that the aging of ASH/ADL may impact AWA/AWC neurons through the intermediary of AMsh glia. We validated this hypothesis by inducing ASH/ADL aging through DAF-16 depletion and found that triggering ASH/ADL aging indeed delayed AMsh glial aging (Figure S2G). These results support the conclusion that the aging of ASH/ADL neurons could suppress AMsh glial aging, thereby protecting AWA/AWC neurons from aging-associated functional decline.

Next, we further investigated how ASH/ADL neuronal aging affected AMsh glia by comparing AMsh glial protein profiles between control and ASH/ADL-aged animals. Adapting a strategy similar to that employed in neurons^36^, we selectively expressed free TurboID in AMsh glia and temporally labeled proteins with biotin only in AMsh glia of adult day 6 animals (Figure 2A). After enrichment by streptavidin-coated beads, the samples were subjected to HPLC-MS/MS quantitative proteomic analyses (Figures 2A, 2B, S3A, and S3B). In both control and experimental groups, we were able to quantitatively characterize approximately 1500 proteins (Table S3), of which 111 were downregulated, and 77 were upregulated in ASH/ADL-aged animals when compared with the same age control animals (Figure 2C). Gene ontology analyses show that the upregulated proteins were predominantly enriched in the endoplasmic reticulum unfolded protein responses (UPR^ER^), specifically the IRE1-XBP1 pathway, suggesting that the aging of ASH/ADL may activate the UPR^ER^ pathways in AMsh glia (Figures 2C and 2D).

**Figure 2.**
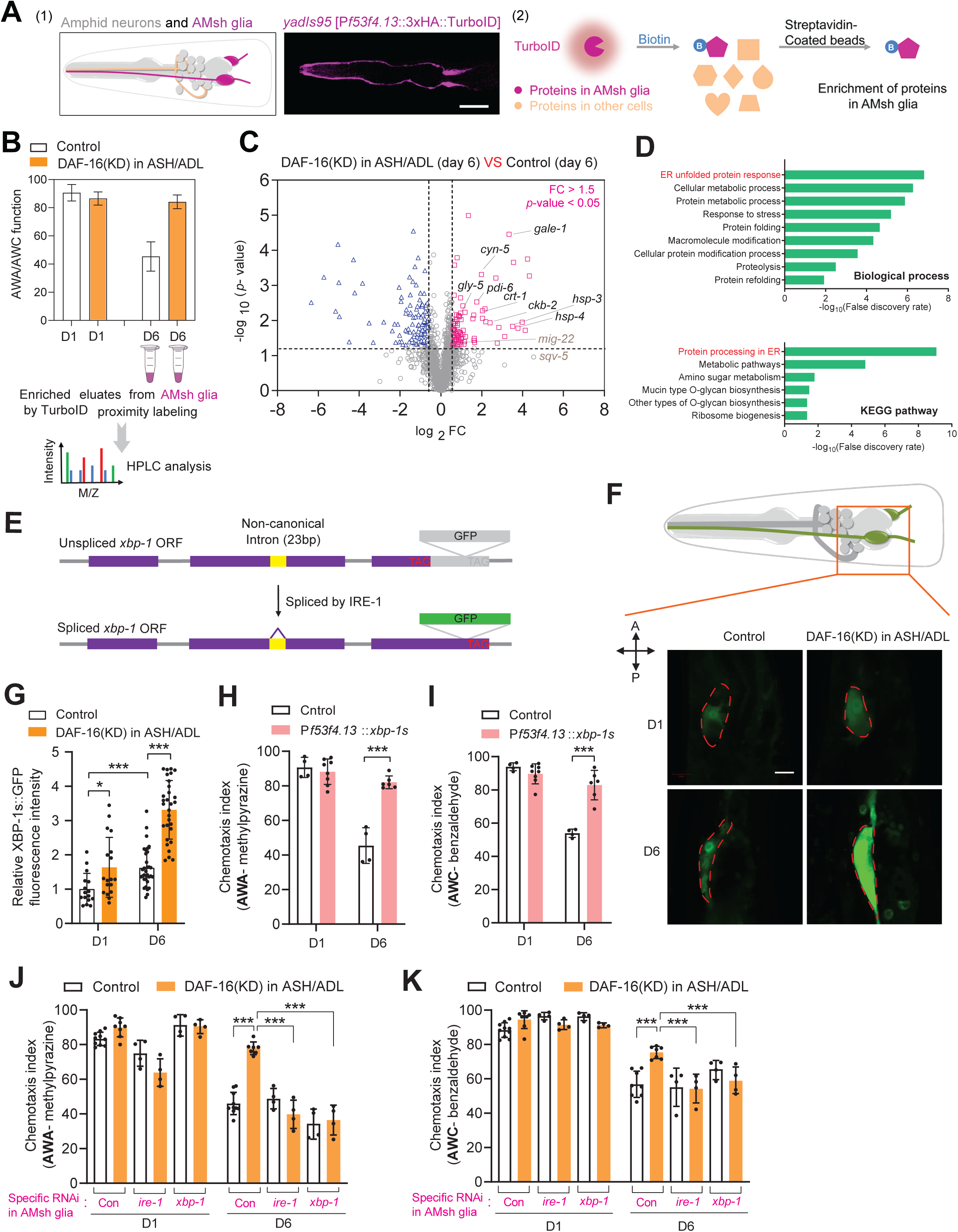
ASH/ADL neuronal aging activates the UPR^ER^ pathway in AMsh glia. **(A)** Schematic of proximity labeling in AMsh glia. (1) An confocal image shows the distribution of HA-tagged TurboID in AMsh glia by immunostaining using anti-HA antibody. Scale bar: 20 μm. (2) A diagram shows the enrichment of biotinylated AMsh glial proteins from worm lysis. **(B)** After proximity labeling, AMsh glial proteins extracted from day 6 control and ASH/ADL DAF-16 depletion (DAF-16 (KD)) animals were subjected to HPLC-MS/MS analyses. **(C)** The Volcano plot of AMsh glial proteomes show proteins with significant changes in DAF-16 (KD) animals when compared with the same age control animals (day 6). log_2_FC > 0.6 or log_2_FC < −0.6 (FC, fold change) with –log_10_(*p-*value) > 1.3 is defined as significantly upregulated (pink square) or downregulated (blue triangle), respectively. The proteins labeled on the plot are selected examples that are known downstream targets of the IRE1-XBP1 pathway. **(D)** Results from gene ontology analyses for proteins that are upregulated by DAF-16 (KD) in ASH/ADL. **(E)** Schematic diagram to illustrate the use of *xbp-1* splicing as a reporter for the activation of the IRE1-XBP1 pathway. In this reporter (*xbp-1s* reporter), *xbp-1*(23bp-intron)::GFP was expressed only in AMsh glia under a AMsh-specific promoter, P*f53f4.13*. GFP was fused at the C-terminal in frame with the spliced version of *xbp-1* mRNA (*xbp-1s*) only when the non-canonical 23bp intron is removed by IRE-1 **(F** and **G)** Confocal images (**F**) and quantifications (**G**) show the expression of the *xbp-1s* reporter in AMsh glia in control and DAF-16(KD) animals at the adult stage day 1 (D1) and day 6 (D6). These experiments were carried out in *daf-16(ot853)* animals with (DAF-16 (KD)) and without (Control) atTIR1 expression in ASH/ADL neurons. The DAF-16::mNG were not expressed in AMsh glia (Figure S1F). ‘A’ indicates the anterior of animals; ‘P’ indicates the posterior of animals. Scale bar: 10μm. Data are shown as mean ± SD. N> 30. One-way ANOVA, *P < 0.05, ***P < 0.001. **(H** and **I)** Results from testing AWA (**H**) and AWC (**I**) neuronal functions by chemotaxis assays using 1% 2-methylpyrazine and 0.5% benzaldehyde, respectively. P*f53f4.13* was used to specifically express the constitutively activated XBP-1s in AMsh glia. Data are shown as mean ± SD. Student’s *t*-test, ***P < 0.001. **(J** and **K)** Results from testing AWA (**J**) and AWC (**K**) neuronal functions in control and DAF-16(KD) animals with and without specifically RNAi knockdown of *ire-1* or *xbp-1* in AMsh glia in day 1 and day 6 animals. Data are shown as mean ± SD. One-way ANOVA, ***P < 0.001.

As one of the major signals mediating the UPR^ER^, the IRE1-XBP1 pathway can regulate the expression of a subset of UPR^ER^ target genes related to protein folding, protein translocation, ER-associated protein degradation, and lipid homeostasis^49–51^. The key event for the activation of the IRE1-XBP1 pathway is the expression of a spliced version of Xbp1(Xbp1s) through the excision of a non-canonical intron from the unspliced Xbp1(Xbp1u) mRNA by the endonuclease IRE1^52^. The spliced XBP1s functions as a transcription factor to regulate the expression of target genes^52^. To examine the potential role of ASH/ADL aging in activating the IRE1-XBP1 pathway in AMsh glia, we expressed the XBP-1 activation reporter in AMsh glia using the similar strategy as previously reported^53^. In this reporter, GFP was fused at the C-terminus in frame with the spliced *xbp-1s* when the non-canonical 23bp intron is removed by IRE-1(Figure 2E). Utilizing this reporter, we can assess the activation of the IRE1-XBP1 pathway by measuring the GFP intensity. Our results showed that the activation of IRE1-XBP1 pathway in AMsh glia was upregulated during aging, and inducing aging in ASH/ADL neurons further enhanced this activation (Figures 2F and 2G). Moreover, we found that expression of the constitutively activated XBP-1s in AMsh glia prevented the functional decline of AWA/AWC neurons during aging (Figures 2H and 2I). Furthermore, AMsh glia-specific knockdown of *xbp-1* or *ire-1*, two key molecules in the IRE1-XBP1 pathway, suppressed the protective effects of ASH/ADL aging on AWA/AWC neurons (Figures 2J and 2K). These results demonstrate that ASH/ADL aging prevents the aging-associated functional decline of AWA/AWC neurons through the activation of the IRE1-XBP1 pathway in AMsh glia.

### Neuronal HSP-4 activates the glial IRE1-XBP1 pathway

Heat shock proteins (HSPs) are well-known components of the UPR^ER^ pathways, and different HSPs have distinct functions, either as downstream factors or upstream regulators of the IRE1-XBP1 pathway^52,54,55^. In the AMsh glial proteome profiling study, we identified two HSPs, HSP-3 and HSP-4, whose expressions in AMsh glia were upregulated by ASH/ADL aging (Figure 2C). To further explore their roles during aging, we first used a *hsp-4* knockin strain, *utx39* [*hsp-4*::mNG::3xFlag], as a reporter to validate our proteomic results and confirmed that the aging of ASH/ADL neurons increased HSP-4 expression in AMsh glia (Figures 3A and 3B). Next, we examined the functional significance of HSP-4 during aging by overexpressing HSP-4 in AMsh glia. In this experiment, we generated transgenes that overexpressed HSP-4 in ASH/ADL neurons as a control. Presumably, HSP-4 would function in AMsh glia but not in ASH/ADL neurons to regulate AWA/AWC functions. Surprisingly, our results showed that the expression of HSP-4 in both AMsh glia and ASH/ADL neurons prevented the aging-associated functional decline of AWA/AWC neurons in a similar fashion as that in *xbp-1s* transgenes (Figures 3C and 3D). To confirm our observations from these gain-of-function studies, we conducted loss-of-function studies by knocking down *hsp-4* in AMsh glia or ASH/ADL neurons. Our results showed that knockdown of *hsp-4* in either AMsh glia or ASH/ADL neurons suppressed the AWA/AWC protections mediated by ASH/ADL-aging (Figures 3E and 3F). Furthermore, knockdown of *hsp-4 s*imultaneously in both AMsh glia and ASH/ADL neurons did not further enhance their suppression on ASH/ADL-aging mediated AWA/AWC protections (Figures 3E and 3F), suggesting that ASH/ADL neuronal and AMsh glial HSP-4 function through the same mechanism. To investigate how neuronal HSP-4 function in this process, we generated HSP-4::mCherry reporters that labels HSP-4 only in ASH/ADL neurons or in all AMsh- channel neurons. Unexpectedly, besides neuronal expression, we also observed HSP- 4::mCherry within AMsh glia in both reporters (Figures 3G and S4A). As a control we expressed free mCherry in all AMsh-channel neurons and didn’t detect any mCherry signal in AMsh glia (Figures 3G and S4A). These results suggest that neuronal HSP-4 is transferred from ASH/ADL neurons to AMsh glia.

**Figure 3.**
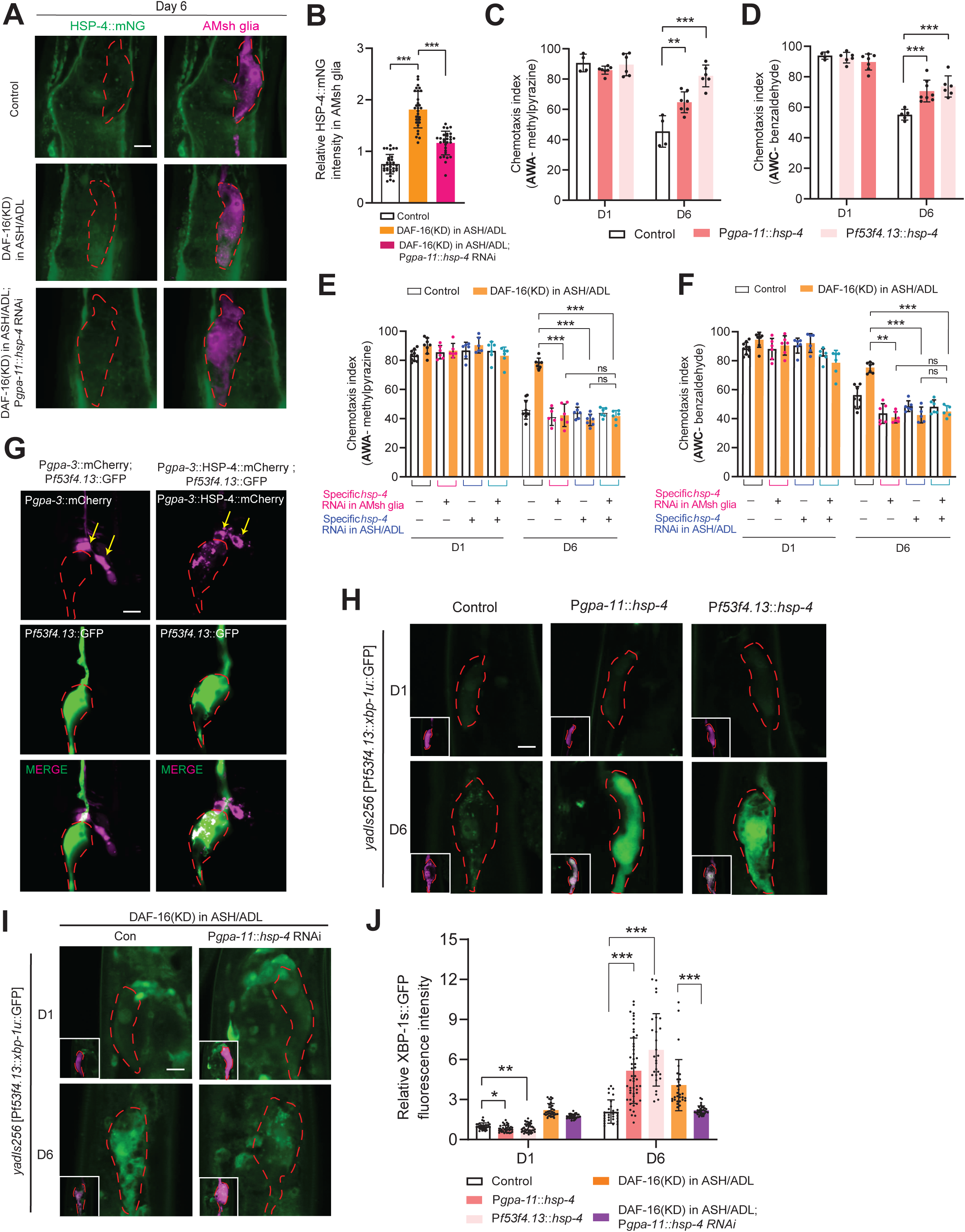
Transmission of HSP-4 from neurons to glia activates the IRE1-XBP1 pathway in AMsh glia. **(A** and **B)** Confocal images (**A**) and quantifications (**B**) show HSP-4::mNG expressions in AMsh glia in control, DAF-16(KD), and DAF-16(KD) with ASH/ADL neuronal *hsp-4* knockdown animals. *gpa-11* promoter was used to drive cell-specific expression in ASH/ADL. *yadIs222* [P*f53f4.13*::mCherry] was used as an AMsh glia marker. Scale bar: 10μm. N>30. Data are shown as mean ± SD. One-way ANOVA, ***P < 0.001 **(C** and **D)** Results from chemotaxis assays show AWA (**C**) and AWC (**D**) neuronal functions in control and transgenes expressing *hsp-4* in ASH/ADL neurons or AMsh glia. *gpa-11* promoter was used to drive cell-specific expression in ASH/ADL neurons. *f53f4.13* promoter was used to drive cell-specific expression in AMsh glia. Data are shown as mean ± SD. One-way ANOVA, **P < 0.01, ***P < 0.001 **(E** and **F)** Results from chemotaxis assays show AWA (**E**) and AWC (**F**) neuronal functions in control, DAF-16(KD), and DAF-16(KD) with *hsp-4* knockdown in AMsh glia, ASH/ADL neurons, or both. *gpa-11* promoter was used to drive cell-specific expression in ASH/ADL neurons. *f53f4.13* promoter was used to drive cell-specific expression in AMsh glia. Data are shown as mean ± SD. One-way ANOVA, **P < 0.01, ***P < 0.001, ns, no significant difference. **(G)** Confocal images show the expression of HSP-4::mCherry or mCherry when they were specifically expressed in AMsh-channel neurons. *gpa-3* promoter was used to drive cell-specific expression in AMsh-channel neurons. *yadIs48* [P*f53f4.13*::GFP] was used as a marker to label AMsh glia. The yellow arrows point to the cell bodies of AMsh-channel neurons. Scale bar: 10 μm. **(H)** Confocal images from the *xbp-1s* reporter (*yadIs256* [P*f53f4.13*::*xbp-1s*(23bp-intron)::GFP]) show the effects of expressing *hsp-4* in ASH/ADL neurons or AMsh glia. *gpa-11* promoter was used to drive cell-specific expression in ASH/ADL neurons. *f53f4.13* promoter was used to drive cell-specific expression in AMsh glia. *yadIs222* [P*f53f4.13*::mCherry] was used as an AMsh glia marker. Scale bar: 10μm. **(I)** Confocal images show the expression of the *xbp-1s* reporter in control and DAF-16(KD) animals with or without knockdown of *hsp-4* in ASH/ADL. *gpa-11* promoter was used to drive cell-specific expression in ASH/ADL neurons. *f53f4.13* promoter was used to drive cell-specific expression in AMsh glia. *yadIs222* [P*f53f4.13*::mCherry] was used as an AMsh glia marker. Scale bar: 10μm. **(J)** Quantification of XBP-1s::GFP fluorescence intensity in **H** and **I**. Data are shown as mean ± SD. N>30. One-way ANOVA, *P < 0.05, **P < 0.01, ***P < 0.001.

As the activation of the IRE1-XBP1 pathway in AMsh glia is regulated by ASH/ADL neuronal aging, we questioned whether ASH/ADL neuronal HSP-4 could be the signal to trigger this activation. To test this, we examined the effects of expressing *hsp-4* in ASH/ADL neurons on the activation of the IRE1-XBP1 pathway in AMsh glia. We observed a significant activation of *xbp-1s* reporter in AMsh glia in aged animals when *hsp-4* was overexpressed in ASH/ADL neurons (Figures 3H and 3J). These results suggested that either HSP-4 itself or some unknown neuronal factors associated with HSP-4 can activate the IRE1-XBP1 pathway in AMsh glia. To distinguish between these possibilities, we tested whether the direct expression of *hsp-4* in AMsh glia could activate the IRE1-XBP1 pathway and found that expressing *hsp-4* in AMsh glia caused the activation of the *xbp-1s* reporter in a similar fashion as that in neuronal transgenes (Figures 3H and 3J). To test whether the role of HSP-4 is unique in neuron-glia communication, we examined the function of HSP-3, another heat shock protein identified in our proteomic studies, in this process. We found that although HSP-3 can be transmitted from AMsh-channel neurons to AMsh glia in a manner similar to HSP-4, overexpression of *hsp-3* in ASH/ADL or AMsh glia didn’t affect the activation of *xbp-1s* reporter in AMsh glia and couldn’t protect AWA/AWC neurons from the aging-associated functional decline (Figures S4B-S4F). These results support the conclusion that the transmission of HSP-4 from ASH/ADL neurons to AMsh glia can activate the IRE1-XBP1 pathway in AMsh glia.

As a homolog of mammalian Hsp70, HSP-4 has been shown to be one of the downstream targets of XBP-1 in unfolded protein responses^52,56^. We tested whether the activation of *xbp-1* could also induce *hsp-4* expression in AMsh glia and found that expression of the constitutively activated XBP-1s increased HSP-4 expression in AMsh glia (Figures S4G and S4H). With these findings showing that HSP-4 acts both upstream and downstream of XBP-1, we believe that the neuronal HSP-4 serves as the initial signal to trigger the activation of the IRE1-XBP1 pathway in AMsh glia, and the activation of XBP-1s induced the expression of glial HSP-4, further enhancing the activation of the IRE1-XBP1 pathway. This positive feedback loop is achieved by an increase in glial HSP-4 due to neuronal HSP-4. To test whether this is the case, using the HSP-4 knockin reporter, we examined the expression of HSP-4 in AMsh glia when HSP-4 was knocked down in ASH/ADL neurons. Consistent with our model, the knockdown of HSP-4 in ASH/ADL neurons suppressed the increase in HSP-4 levels in AMsh glia caused by ASH/ADL neuronal aging (Figures 3A and 3B). The essential role of neuronal HSP-4 in this neuron-glia interaction was further confirmed by the behavior analyses and *xbp-1s* reporter; the knockdown of *hsp-4* in ASH/ADL neurons attenuated the protection of AWA/AWC neurons and the activation of *xbp-1s* reporter caused by ASH/ADL neuronal aging (Figures 3E, 3F, 3I, and 3J). Collectively, our data show that the transmission of HSP-4 from aged ASH/ADL neurons to AMsh glia can activate the glial IRE1-XBP1 pathway.

### Neuronal HSP-4 is transmitted to AMsh glia through extracellular vesicles

With the findings of the function of neuronal HSP-4 in activating the glial IRE1-XBP1 pathway, we further asked what could mediate the transmission of HSP-4 from neurons to glia. Neuronal HSP-4::mCherry displayed both puncta and diffused distributions in AMsh glia, and these two different patterns may represent two stages of neuronal HSP-4 in AMsh glia: the early stage when it is still packaged in vesicles from neurons, and the late stage when it is incorporated into AMsh glia (Figure 4A). To understand the machinery that mediates the transmission of HSP-4 from neurons to glia, we focused our analyses on the HSP-4::mCherry puncta in AMsh glia. After examining markers for different neuronal vesicles, we found that the TSP-6::GFP-labeled neuronal extracellular vesicles (EV) were uptake by AMsh glia^57,58^, and the HSP-4::mCherry puncta within AMsh glia are highly co-localized with these neuronal EVs (Figures 4A, S5A, and S5B). Consistent with our findings, HSP-4 and its homologs were found to be packaged in EVs released from neurons in *C. elegans* and mammals^58,59^.

**Figure 4.**
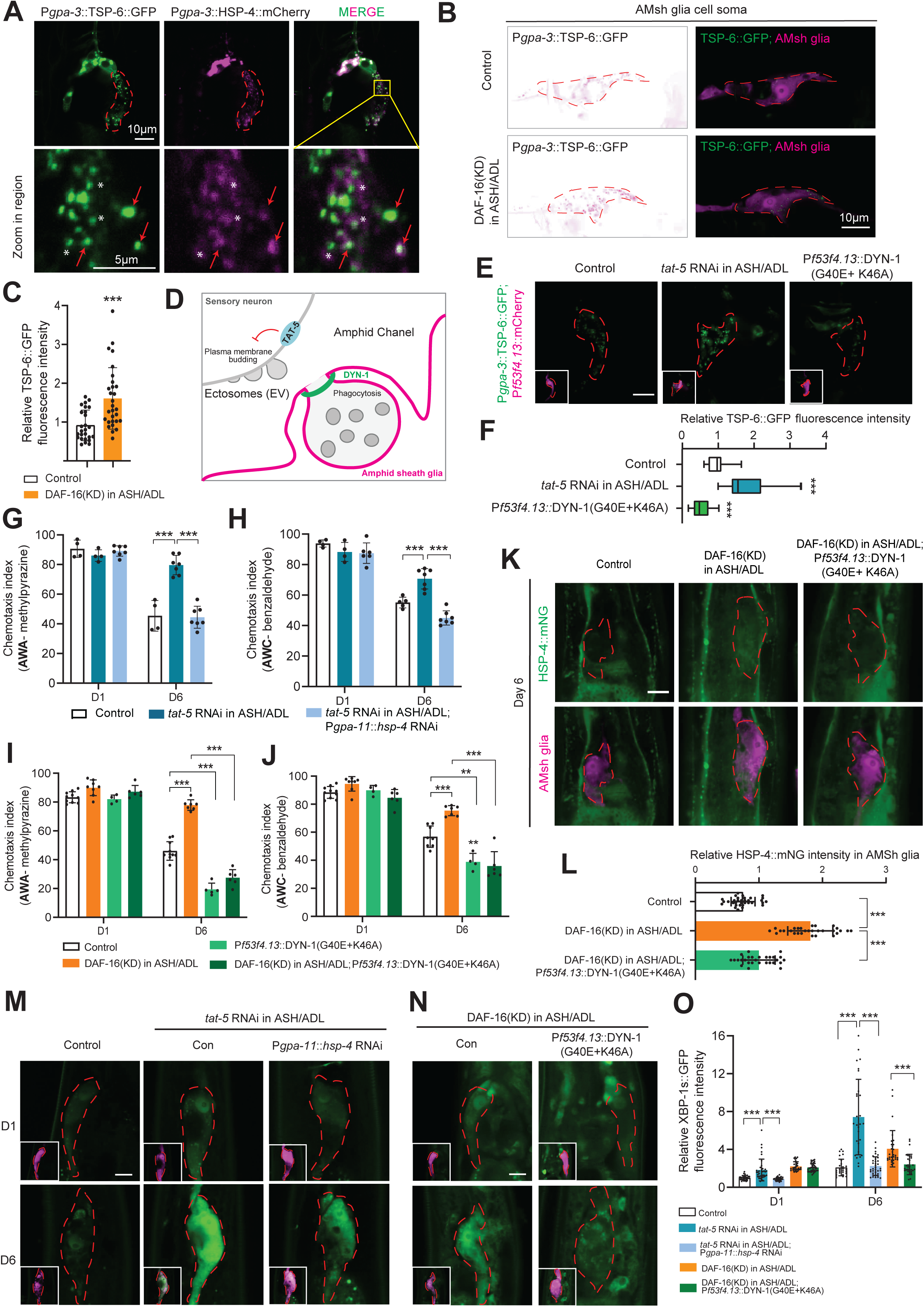
Neuronal HSPs are transmitted to AMsh glia through extracellular vesicles. **(A)** Confocal images show that neuronal HSP-4::mCherry is co-localized with TSP-6::GFP, a extracellular vesicle marker expressed in AMsh-channel neurons, in AMsh glia. *gpa-3* promoter was used to drive cell-specific expression in AMsh-channel neurons. The red arrows point to the co-localized HSP-4::mCherry and TSP-6::GFP puncta. The white asterisks indicate the diffused HSP-4::mCherry. In the upper panel, scale bar, 10μm. In the zoom in region, scale bar, 5μm. **(B** and **C)** Confocal images (**B**) and quantifications (**C**) show the signal from neuronal expressed TSP-6::GFP in AMsh glia in control and DAF-16(KD) animals. TSP-6::GFP expression was driven by the AMsh-channel neurons specific promoter *gpa-3 (yadIs226* [P*gpa-3*::*tsp-6*::GFP]). *yadIs222* [P*f53f4.13*::mCherry] was used as an AMsh glia marker. Scale bar: 10μm. N>30. Data are shown as mean ± SD. Student’s *t*-test, ***P < 0.001. **(D)** Schematic diagram shows that the release and uptake of extracellular vesicles (EV) are regulated by TAT-5 and DYN-1, respectively. **(E** and **F)** Confocal images (**E**) and quantifications (**F**) of *yadIs226* [P*gpa-3*::*tsp-6*::GFP] signals in control, knockdown of *tat-5* in ASH/ADL, and expression of the dominant-negative DYN-1(G40E+K46A) in AMsh glia. *gpa-3* promoter was used to drive cell-specific expression in AMsh-channel neurons. *gpa-11* promoter was used to drive cell-specific expression in ASH/ADL neurons. *f53f4.13* promoter was used to drive cell-specific expression in AMsh glia. *yadIs222* [P*f53f4.13*::mCherry] was used as an AMsh glia marker. Scale bar: 10μm. N>30. Data are shown as mean ± SD. One-way ANOVA, ***P < 0.001. **(G** and **H)** Results from chemotaxis assays show AWA (**G**) and AWC (**H**) neuronal functions in control, ASH/ADL *tat-5* knockdown, and ASH/ADL *tat-5*&*hsp-4* knockdown animals. *gpa-11* promoter was used to drive cell-specific expression in ASH/ADL neurons. Data are shown as mean ± SD. One-way ANOVA ***P < 0.001. **(I** and **J)** Results from chemotaxis assays show AWA (**I**) and AWC (**J**) neuronal functions in control and DAF-16(KD) animals with or without expressing the dominant-negative DYN-1(G40E+K46A) in AMsh glia. Data are shown as mean ± SD. One-way ANOVA, **P < 0.01, ***P < 0.001. **(K** and **L)** Confocal images (**K**) and quantifications (**L**) show HSP-4::mNG expression in AMsh glia in control and DAF-16(KD) animals with and without expression of the dominant-negative DYN-1(G40E+K46A) in AMsh glia. *gpa-11* promoter was used to drive cell-specific expression in ASH/ADL neurons. *f53f4.13* promoter was used to drive cell-specific expression in AMsh glia. *yadIs222* [P*f53f4.13*::mCherry] was used as an AMsh glia marker. Scale bar:10μm. N>30. Data are shown as mean ± SD. One-way ANOVA, ***P < 0.001. **(M)** Confocal images show the expression of the *xbp-1s* reporter, *yadIs256* [P*f53f4.13*::*xbp-1s*(23bp-intron)::GFP], in animals with or without *tat-5* knockdown in ASH/ADL neurons and in animals with knockdown of *tat-5* and *hsp-4* simultaneously in ASH/ADL neurons. *gpa-11* promoter was used to drive cell-specific expression in ASH/ADL neurons. *f53f4.13* promoter was used to drive cell-specific expression in AMsh glia. *yadIs222* [P*f53f4.13*::mCherry] was used as an AMsh glia marker. Scale bar: 10μm. **(N)** Confocal images show the expression of the *xbp-1s* reporter, *yadIs256* [P*f53f4.13*::*xbp-1s*(23bp-intron)::GFP], in DAF-16(KD) animals with and without expression of the dominant-negative DYN-1(G40E+K46A) in AMsh glia. *gpa-11* promoter was used to drive cell-specific expression in ASH/ADL neurons. *f53f4.13* promoter was used to drive cell-specific expression in AMsh glia. *yadIs222* [P*f53f4.13*::mCherry] was used as an AMsh glia marker. Scale bar: 10μm. **(O)** Quantifications of XBP-1s::GFP fluorescence intensity in **M** and **N**. N>30. Data are shown as mean ± SD. One-way ANOVA, ***P < 0.001.

Next, we examined the effects of ASH/ADL neuronal aging on EVs and observed more neuronal EVs in AMsh glia in ASH/ADL-aged animals when compared with the same age control animals, supporting that neuronal aging can lead to more EVs being transmitted from neurons to glia (Figures 4B and 4C). To further elucidate the role of neuronal EVs in regulating AMsh glia, we genetically manipulated EV release and uptake and examined the consequences of these modulations. In *C. elegans*, *tat-5,* a phospholipid flippase, inhibits the budding of EVs from the plasma membrane^60,61^, and *dyn-1*, a homolog of mammalian dynamin 2, is required for EVs uptake is glial cells^57,62,63^ (Figure 4D). We first confirmed the function of *tat-5* and *dyn-1* in our system, that knockdown of *tat-5* in AMsh-channel neurons increased EV release, whereas expression of a dominant-negative DYN-1(G40E and K46A)^57,63^ in AMsh glia suppressed EV uptake (Figures 4E and 4F). We then examined the effects of increasing neuronal EV release on the activation of the IRE1-XBP1 pathway in AMsh glia and on the protection of AWA/AWC function by AMsh glia. We found that increasing EV release from ASH/ADL neurons by cell specific knockdown of *tat-5* upregulated the extent of IRE1-XBP1 activation and prevent the aging-associated AWA/AWC functional decline (Figures 4G, 4H, 4M, and 4O). More importantly, the function of EVs depended on neuronal HSP-4, as the knockdown of *hsp-4* in ASH/ADL neurons abrogated the effects of increasing EV release (Figures 4G, 4H, 4M, and 4O). Furthermore, we found that the suppression of EV uptake through overexpressing the dominant-negative DYN-1 in AMsh glia attenuated the upregulation of HSP-4 proteins and the activation of the IRE1-XBP1 pathway in AMsh glia in ASH/ADL-aged animals (Figures 4K, 4L, 4N, and 4O). Moreover, suppressing AMsh glial EV uptakes largely eliminated the protections of AWA/AWC functions by ASH/ADL neuronal aging (Figures 4I and 4J). Together, our results support the conclusion that ASH/ADL neuronal HSP-4 is transmitted to AMsh glia through EVs, and this transmission activates the glial IRE1-XBP1 pathway to protect AWA/AWC neurons from functional decline during aging.

### AMsh glial chondroitin protects neurons from aging

With all these findings, we sought to investigate how the activation of the IRE1-XBP1 pathway in AMsh glia protected AWA/AWC neurons from aging-associate functional decline. To address this question, we focused on proteins that are upregulated by ASH/ADL aging and tested the effects of overexpressing them in AMsh glia on AWA/AWC functions (Figure 5A). We found that overexpression of *mig-22,* a chondroitin polymerizing factor and a homolog of mammalian *ChSy2*^64,65^, suppressed the functional decline of AWA/AWC neurons during aging (Figures 5B and 5C). This data suggests that the increase in chondroitin synthesis could be the downstream signal triggered by the activation of the IRE1-XBP1 pathway, mediating the protection of AWA/AWC neurons against aging. Since the expression of the only chondroitin synthase in *C. elegans*^66,67^, *sqv-5*, is also highly upregulated in AMsh glia by ASH/ADL aging (Figure 2C; Table S3), we tested the function of *sqv-5* in AWA/AWC protections and found that overexpression of *sqv-5* in AMsh glia suppressed the aging-associated AWA/AWC functional decline in a similar manner as that in *mig-22* transgenes (Figures 5B and 5C). Using MIG-22::GFP and SQV-5::GFP translational reporters driven by their own promoters, we confirmed that their expressions in AMsh glia were upregulated by ASH/ADL aging, and these increases depended on the activation of the IRE1-XBP1 pathway in AMsh glia (Figures 5D-5G). In addition, we found that the expression of the constitutively activated XBP-1s in AMsh glia is sufficient to drive MIG-22::GFP and SQV-5::GFP expression in AMsh glia (Figures 5H-5J). Furthermore, AMsh glia-specific knockdown of *mig-22* or *sqv-5* suppressed the protections of AWA/AWC function by ASH/ADL neuronal aging (Figures 5K and 5L). Collectively, these results highlighted the important roles of glia-produced chondroitin in the protection of AWA/AWC neurons during aging.

**Figure 5.**
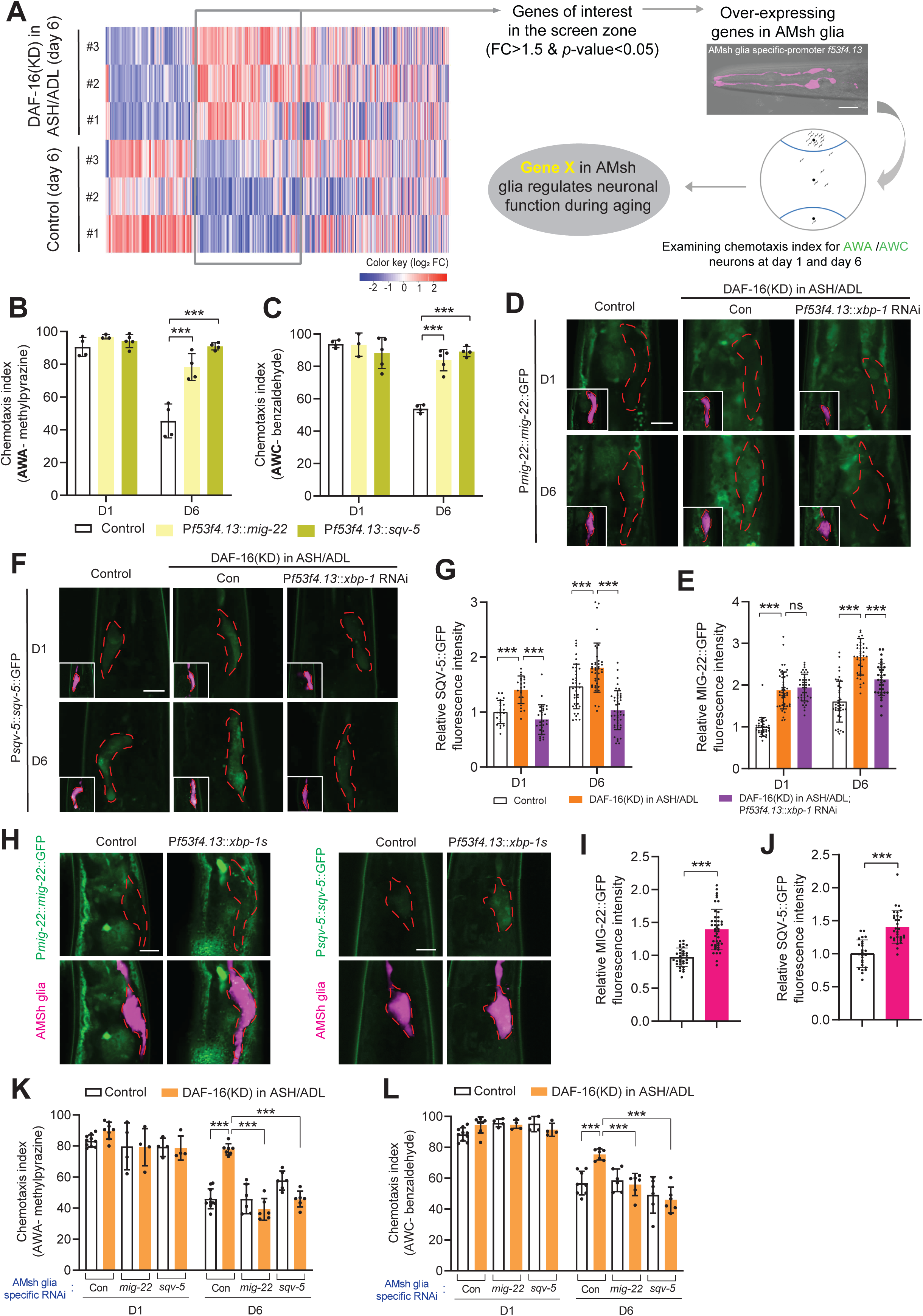
Expression of chondroitin in AMsh glia protects AWA/AWC neurons from aging-associated functional decline. **(A)** A strategy for testing the function of DAF-16 depletion-upregulated genes in AWA/AWC functions. The genes of interest were overexpressed in AMsh glia to test their effects on AWA/AWC neuronal functions of day 1 and day 6 animals. **(B** and **C)** Results from chemotaxis assays show AWA (**B**) and AWC (**C**) neuronal functions in control and transgenes expressing *mig-22* or *sqv-5* in AMsh glia. *f53f4.13* promoter was used to drive cell-specific expression in AMsh glia. Data are shown as mean ± SD. One-way ANOVA, ***P < 0.001 **(D** and **E)** Confocal images (**D**) and quantifications (**E**) show the expression of MIG-22::GFP in AMsh glia in control and DAF-16(KD) animals with or without *xbp-1* RNAi knockdown in AMsh glia. *yadIs272* [P*mig-22*::*mig-22*::GFP] is a translational reporter for *mig-22*. *yadIs222* [P*f53f4.13*::mCherry] was used as an AMsh glia marker. Scale bar, 10μm. N>30. Data are shown as mean ± SD. One-way ANOVA, ***P < 0.001, ns, no significant difference **(F** and **G)** Confocal images (**F**) and quantifications (**G**) show the expression of SQV-5::GFP expression in AMsh glia in control and DAF-16(KD) animals with or without *xbp-1* RNAi knockdown in AMsh glia. *yadCK287* [P*sqv-5*::*sqv-5*::GFP] is a knockin reporter for *sqv-5* expression. *yadIs222* [P*f53f4.13*::mCherry] was used as an AMsh glia marker. Scale bar: 10μm. N>30. Data are shown as mean ± SD. One-way ANOVA, ***P < 0.001. **(H** to **J)** Confocal images (**H**) and quantifications show the expression of MIG-22::GFP (**I**) and SQV-5::GFP (**J**) in animals with or without expressing constitutively activated *xbp-1s* in AMsh glia. *f53f4.13* promoter was used to express the constitutively activated *xbp-1s* in AMsh glia. *yadIs222* [P*f53f4.13*::mCherry] was used as an AMsh glia marker. Scale bar: 10μm. N>30. Data are shown as mean ± SD. Student’s *t*-test, ***P < 0.001. **(K** and **L)** Results from chemotaxis assays show AWA (**K**) and AWC (**L**) neuronal functions in control and DAF-16(KD) animals with or without specifically knockdown of *mig-22* or *sqv-5* in AMsh glia. Data are shown as mean ± SD. One-way ANOVA, ***P < 0.001.

## Discussion

In conclusion, we have characterized a distinctive mechanism of neuron-glia communication during aging, wherein glia sense and respond to early-aged neurons through released neuronal extracellular vesicles (EVs). The neuronal HSP-4 packaged in EVs serves as a signal transmitted into glia, activating the glial UPR^ER^ pathway in response to neuronal aging. Furthermore, the activated UPR^ER^ pathway in glia transcriptionally regulates chondroitin synthases, thereby providing protection against neuronal aging (Figure S6). Given the ubiquity of HSPs, UPR^ER^ and EVs across species, the mechanism we have elucidated may represent a potentially conserved process within the nervous system during aging.

In the brain, different neurons respond to stimulations differently in physiological and pathological conditions, with some neurons being more sensitive than others. In this study, we demonstrate that early-aged neurons can induce cellular responses in glia through EV- mediated signal transductions, thereby protecting other neurons from aging-associated functional decline. This mechanism may represent a more general process underlying neuronal protection beyond aging: when stress or damage occurs, a subset of neurons can respond to the stimulations to activate surrounding glia through EVs, and the activation of glial can, in turn, protects other neurons and prevents cascade damage to the brain. Consistent with this idea, under stress or after damage, many cells, including neurons, can release EVs, and those EVs contain a wide array of constituents, such as proteins, DNA, RNA, lipids and metabolites^15^. Although this study primarily focuses on the role of EV- packaged HSPs in neuronal protection, we believe that many components of EVs could play important roles in signal transduction within the nervous system. Since EVs can mediate both short- and long-distance intercellular communication^15^, the mechanism presented in this study may also provide insights in many biological processes that are initiated by neurons. For instance, in *C. elegans*, activation of stress responses in neurons and glia can affect lifespan through intestinal cells, and the mechanisms mediating this cell-nonautonomous signal transduction from the brain to peripheral tissues remain unclear^52,68–70^. It is possible that neurons and glia transmit signaling molecules to intestinal cells through EVs to affect the overall heathy of animals.

Many physiological and pathological factors, including aging, can lead to ER stress^71–74^. To maintain intracellular homeostasis, cells have evolved an adaptive mechanism, unfolded protein response of the ER (UPR^ER^), to mitigate the ER stress^51^. The activation of UPR^ER^ can be regulated by cell-autonomous and cell-nonautonomous mechanisms^52,70^. In this study, we investigate the cell-nonautonomous regulation of UPR^ER^ between neuron and glia, and we show that the activation of glial UPR^ER^ can be initiated by signals from neurons, highlighting the significant role of neurons in regulating glial homeostasis during aging. As one of the major UPR^ER^ signals, the IRE1-XBP1 pathway control the expression of many target genes, including Hsp70, and Hsp70 can suppress the IRE1-XBP-1 activation in cultured cells^71,77,78^. We found that *C. elegans* HSP-4, a homolog of mammalian Hsp70, can be transmitted from neurons to glia during aging, and neuronal HSP-4 triggers the activation of the glial IRE1-XBP1 pathway during aging. The differences of mammalian Hsp70 and *C. elegans* HSP-4 in the activation of the IRE1-XBP1 pathway are likely caused by aging, as we found that expression of HSP-4 in young adults suppressed the activation of the IRE1-XBP1 pathway in a similar fashion as that in cultured cells (Figures 3H and 3J). Therefore, we believe that the function of HSP-4 in UPR^ER^ activation in our study is attributed to aging-associated changes.

Chondroitin, a type of glycosaminoglycan (GAG), is a linear polysaccharide composed of repeating disaccharide units. Typically, one or more chondroitin chains are covalently bound to core proteins through a common tetra-saccharide linkage region, forming proteoglycans that are secreted into the extracellular matrix^79^. The properties of chondroitin enable it to interact with water, ions, specific chemicals or proteins, maintaining the specific structure of lumens or channels, generating osmotic pressure on its surroundings, and facilitating local signaling within the microenvironment. Thus, chondroitin proteoglycans, as the extracellular matrix component, are involved in intercellular signaling and structure scaffold for the extracellular space, which is required for cytokinesis, embryogenesis, epithelial morphogenesis and cell division^65–67^. Here, we found that the activation of the glial UPR^ER^ pathway can increase the expression of chondroitin synthases, *sqv-5/ChSy1* and *mig-22/ChSy2,* to suppress neuronal aging, supporting the critical roles of glial chondroitin in neuronal protection. As the sensory cilia of glia-embedded neurons in *C. elegans* are fully wrapped by glia, it is plausible that the chondroitin proteoglycans secreted from glia can provide support for the functional structure of cilia and maintain the microenvironment homeostasis for sensory neurons during aging.

## Acknowledgements

We thank all Yan lab members for comments on the manuscript and double-blind experiments. We are grateful to Dr. John E. Cronan for providing the biotin auxotrophic E. coli strain (MG1655). Randy Cockerell, Alp Altinbasak and Amber White prepared most of the reagents used in this study. Greg Waitt and Tricia Ho contributed to sample preparation and data acquisition within the Duke Proteomics and Metabolomics Core Facility. Some strains were provided by the CGC, which is funded by the NIH Office of Research Infrastructure Programs (P40 OD010440). This project is supported by NIH grants NS105638 and AG073994 (to D.Y.).

## Author Contributions

J.W. and D.Y. conceived the study and interpreted all the data. J.W. performed all molecular, genetic, imaging and proteomic experiments. E.S. carried out HPLC-MS and proteomic data analysis. O.Y. performed RNAi experiments. J.W. and D.Y. wrote the manuscript with inputs from all authors.

## Competing financial interests

The authors declare no competing financial interests.

## Methods

### Lead Contact

Further information and requests for resources should be directed to and will be fulfilled by the lead contact, Dong Yan.

### Materials Availability

*C. elegans* strains and plasmids generated in this study are available from the lead contact without restriction.

### *C. elegans* Strains and Maintenance

*C. elegans* were maintained on nematode growth media (NGM) with E. coli OP50 at 20°C, as previously described^80^. Bristol N2 was used as the wild-type strain in this study. For the TurboID proximity labeling experiments, worms were fed on biotin auxotrophic *E. coli* (MG1655 bioB: kan) to eliminate excessive biotin to reduce the background for the AMsh glial proteome. Transgenic strains were produced by injecting plasmids DNA mixes into the hermaphrodite gonad. P*unc-122*::GFP, P*unc-122*::RFP, P*ttx-3*::GFP or P*ttx-3*::RFP served as co-injection markers to facilitate the selection of transgenic animals. Integrated transgenic strains were generated using UV/Trioxsalen treatment. The CRISPR/Cas-9 knockin strain *yadCK287* [P*sqv-5*::*sqv-5*::GFP] was created using a previously established method^81^. All strains used in this study are listed in Table S1.

### Aging related experiments

To synchronize developmental stages for aging-related experiments in *C. elegans*, worm eggs were isolated by treating adult worms with bleach buffer (4 ml of bleach 4%, 1.5 ml 5 M NaOH, 4 ml dH_2_O) and then synchronized at the L1 stage overnight in the absence of food. These synchronized L1 larvae were raised on NGM seeded with OP50 at 20 °C until L4 larvae stage. L4 stage worms were shifted to NGM plates containing OP50 supplemented with 100 μg ml^−1^ 5-Fluoro-2’-deoxyuridine (FUdR) to inhibit progeny development.

### Constructs cloning for expression and RNAi

All constructs used in this study are described in Table S2. The plasmids prepared for microinjection were constructed using the Gateway Cloning technology. DNA fragments from genes or their associated promoters were amplified via polymerase chain reaction (PCR). These fragments were subsequently inserted into the pCR8 entry vector (Invitrogen) or destination vector through Gibson ligation. The expression plasmids were generated through the LR reaction (Invitrogen) by using entry vector and destination vector. For *sqv-5* CRISPR/Cas-9 knockin, the single-guide RNA (sgRNA) was designed following the sequence characterized by G/A(N)19NGG, which was placed into the pU6::G/A(N)19_sgRNA plasmid by overlap extension PCR.

RNAi knockdown specific to AMsh glia, AMsh channel-neurons, and ASH/ADL neurons was performed as previously described^82,83^. For each gene RNAi construct, a 500-1000 bp cDNA fragment was amplified by the polymerase chain reaction (PCR). The purified DNA fragment was then inserted on entry vector pCR8 (Invitrogen). The orientation (sense and antisense) of these DNA fragments in the pCR8 vector was identified through sequencing. Subsequently, they were assembled under various promoters using the LR reaction (Invitrogen) to produce the specific RNAi constructs for each gene. To generate transgenic animals with the target gene silenced in designated cells, the sense and antisense RNAi constructs were mixed with co-injection marker and injected into the gonad of animals.

### TurboID Proximity labeling in Amsh glia

To acquire the proteome in AMsh glia, we utilized a method previously developed in our lab^36^. We generated *yadIs95* [P*f53f4.13*::3×HA::TurboID] to exclusively express free TurboID in Amsh glia of *C. elegans*. For biotin labeling experiments, worms were raised on biotin auxotrophic *E. coli* (MG1655) as food to reduce background biotin levels. Synchronized L1 larval were grown on NGM until L4 stage at 20°C. Subsequently, L4 stage animals were transferred on the NGM plates supplemented with FUdR (5-Fluoro-2’-deoxyuridine, 100μg mL^-1^) and maintained at 20°C. On day 6, 100μM exogenous biotin was added to the FUdR plate and the worms were treated for 8 hours at 25°C. Afterward, these day 6 animals were harvested and washed 3 times with M9 buffer on ice. Finally, worm pellet was resuspended in NP40 lysis buffer for sonication and the lysates were centrifuged at 10000×g at 4°C to remove the debris and lipids. The clarified lysates were used for enrichment by streptavidin-coated beads and for western blot analysis.

To enrich biotinylated proteins from AMsh glia, we first prepared 50 µL of streptavidin-coated magnetic beads by washing them twice with NP40 lysis buffer. These beads were then incubated with the clarified lysates, rotating overnight at 4°C. Next, the beads were washed twice with 1 mL of NP40 lysis buffer, once with 1 mL of 1 M KCl, once with 1 mL of 0.1 M Na_2_CO_3_, once with 1 mL of 2 M urea in 10 mM Tris-HCl (pH 8.0) and twice with 1 mL NP40 lysis buffer. Subsequently, these beads were suspended in 100 µL elution buffer (2% SDS, 25 mM Tris-HCl, 50 mM NaCl, 2 mM biotin, 20 mM DTT). Finally, biotinylated proteins from AMsh glia were eluted in the buffer by boiling the beads at 80°C for 10 min. The eluates were subjected to western blotting and HPLC-MS/MS quantitative proteomic analysis.

### Western blot analysis

Worm lysates were prepared by sonication, followed by centrifugation at 10000×g at 4°C to eliminate the debris. The clarified protein samples mixed with an equivalent volume of protein sample buffer and subsequently denatured at 95°C for 10 min. The denatured protein samples were subjected to electrophoresis using a 4-20% SDS-PAGE gel (Bio-Rad) in running buffer and subsequently transferred onto polyvinylidene difluoride (PVDF) membranes in transferring buffer. Primary antibodies used in this study were rabbit anti-HA antibody (Sigma, H6908) and rabbit anti-Actin antibody (Sigma, A2066) at 1:1000 dilution. The secondary antibody was goat anti-rabbit IgG at 1:5000 dilution (GE Healthcare). Biotinylated proteins were blotted using HRP-conjugated Streptavidin at 1:1000 dilution (Thermo, N100). Images were acquired using a chemiluminescence imaging system (Syngene).

### LC-MS/MS analysis

Quantitative LC/MS/MS was performed on 3 µL using an MClass UPLC system (Waters Corp) coupled to a Thermo Orbitrap Fusion Lumos high resolution accurate mass tandem mass spectrometer (Thermo) equipped with a FAIMSPro device via a nanoelectrospray ionization source. Briefly, the sample was first trapped on a Symmetry C18 20 mm × 180 µm trapping column (5 μl/min at 99.9/0.1 v/v water/acetonitrile), after which the analytical separation was performed using a 1.8 µm Acquity HSS T3 C18 75 µm × 250 mm column (Waters Corp.) with a 90-min linear gradient of 5 to 30% acetonitrile with 0.1% formic acid at a flow rate of 400 nanoliters/minute (nL/min) with a column temperature of 55C. Data collection on the Fusion Lumos mass spectrometer was performed for three difference compensation voltages (−40v, −60v, −80v). Within each CV, a data-dependent acquisition (DDA) mode of acquisition with a r=120,000 (@ m/z 200) full MS scan from m/z 375 – 1500 with a target AGC value of 4e5 ions was performed. MS/MS scans were acquired in the ion trap in Rapid mode with a target AGC value of 1e4 and max fill time of 35 ms. The total cycle time for each CV was 0.66s, with total cycle times of 2 sec between like full MS scans. A 20s dynamic exclusion was employed to increase depth of coverage. The total analysis cycle time for each injection was approximately 2 hours.

### Quantitative MS/MS Data Analysis

Following UPLC-MS/MS analyses, data were imported into Proteome Discoverer 3.0 (Thermo Scientific Inc.). In addition to quantitative signal extraction, the MS/MS data was searched against the SwissProt C. elegans database (downloaded in Aug 2022) and a common contaminant/spiked protein database (bovine albumin, bovine casein, yeast ADH, human keratin, etc.), and an equal number of reversed-sequence “decoys” for false discovery rate determination. Sequest with Infernys enabled (v 3.0, Thermo PD) was utilized to produce fragment ion spectra and to perform the database searches. Database search parameters included fixed modification on Cys (carbamidomethyl) and variable modification on Met (oxidation). Search tolerances were 2ppm precursor and 0.8Da product ion with full trypsin enzyme rules. Peptide Validator and Protein FDR Validator nodes in Proteome Discoverer were used to annotate the data at a maximum 1% protein false discovery rate based on q-value calculations. Note that peptide homology was addressed by only using unique peptides for quantitation. Protein homology was addressed by grouping proteins that had the same set of peptides to account for their identification. A master protein within a group was assigned based on % coverage. Prior to normalization, a filter was applied such that a peptide was removed if it was not measured in at least 50% of the samples in a single group. After that filter, samples were total intensity normalized (total intensity of all peptides for a sample are summed then normalized across all samples). Next, the following imputation strategy is applied to missing values. If less than half of the values are missing in a biological group, values are imputed with an intensity derived from a normal distribution of all values defined by measured values within the same intensity range (20 bins). If greater than half values are missing for a peptide in a group and a peptide intensity is > 5e6, then it was concluded that peptide was misaligned and its measured intensity is set to 0. Peptide intensities were then subjected to a trimmed-mean normalization in which the top and bottom 10 percent of the signals were excluded, and the average of the remaining values was used to normalize across all samples. Lastly, all peptides belonging to the same protein were then summed into a single intensity. These protein level intensities are what were used for the remained of the analysis (Table S3).

### Chemotaxis assay for sensory neurons

Chemotaxis assay for AWA, AWC and ASH/ADL were performed as previously described ^33,38,84^. We labeled the chemotaxis plates (2% agar, 5mM KPO4, 1 mM CaCl_2_, 1 mM MgSO_4_) as shown in Figures S1A and S1B. We added 2 µL of 0.5 M sodium azide to both odorant spots and control spots on all chemotaxis plates. Once the sodium azide was absorbed into the agar, worms were rinsed off NGM plates using M9 buffer and transferred to 1.5 mL centrifuge tube.

Leave the tubes on the bench for 1 min to let the worms settle at the bottom of the tube and remove the supernatant M9 buffer. Next, worms were washed twice with 1 mL M9 buffer, once with chemotaxis assay buffer (5mM KPO4, 1 mM CaCl_2_, 1 mM MgSO_4_). When worms are ready, 80-100 worms suspended in the chemotaxis assay buffer were dropped at the center spot of chemotaxis plates. Subsequently, we added 1.5 µL of ethanol to the control spot and 1.5 µL of designated experimental chemical (1% 2-mythylpyrazine, 0.5% benzaldehyde, or 100% octanol) to the odorant spot. The excess liquid from the worm drop were removed using filter paper.

Animals will move on the surface of the chemotaxis plates to respond to the odorant. The chemotaxis index was calculated as follows:

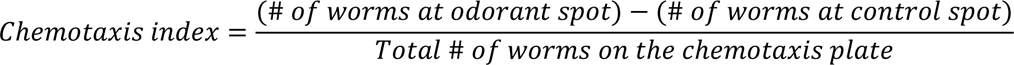

### Auxin-inducible degradation of DAF-16

The Auxin-inducible degradation (AID) system was employed to degrading DAF-16 in specific neurons as described previously^42,43^. *daf-16(ot853)* [*daf-16*::linker::mNeonGreen::3xFlag::AID] was a strain in which a degron-tagged mNeonGreen was inserted at the C-terminus of endogenous *daf-16*. To exclusively deplete DAF-16 in the ASH/ADL neurons, we generate a transgenic line that expresses <u>at</u>TIR1 (derive from *<u>A</u>rabidopsis thaliana*) by promoter *gpa-11* in *daf-16(ot853)* animals. *daf-16(ot853)* without atTIR1 expression was used as control. Transgenic animals were fed on the NGM plates supplemented with 5 μM auxin (indole-3 acetic acid, sigma) during aging.

### Visualization and quantification of fluorescence

Worms were picked and mounted on a 4% agarose pad and treated with 5 mM levamisole. The fluorescence was visualized using a ZEISS fluorescence microscope and ZEISS LSM700 confocal microscope. The fluorescence intensity was quantified by using the Fiji (ImageJ).

### Statistical analysis

Data are presented as the means ± SD unless specifically indicated. Statistical analyses included student’s *t*-test, One-way ANOVA and Two-way ANOVA. All figures were generated using GraphPad Prism 9 (GraphPad Software, La Jolla, CA, USA) and Adobe Illustrator.

## Data availability

The authors declare that all data supporting the findings of this study are available within the article and its Supplementary materials.

## Supplemental information

**Figure S1.**
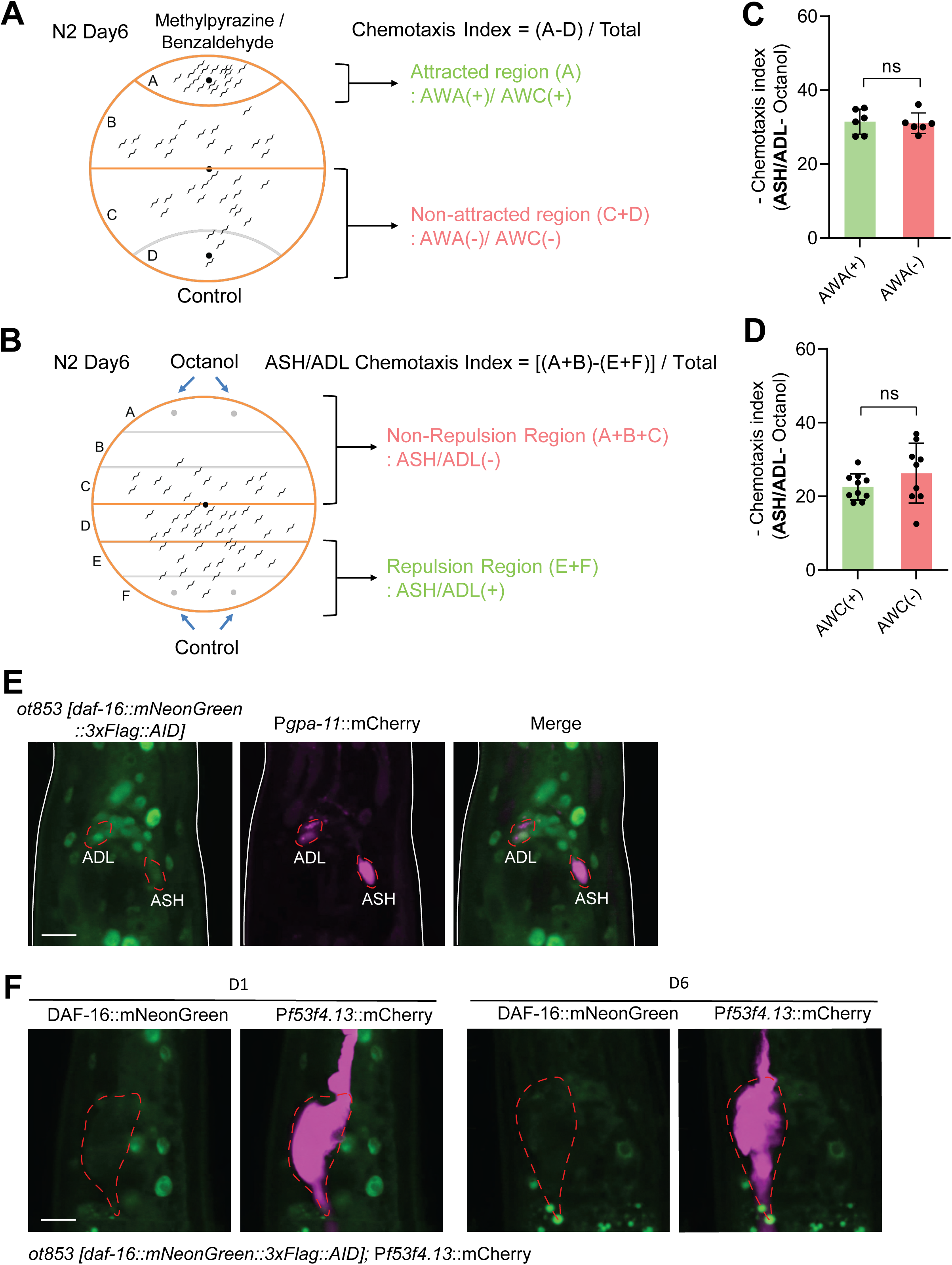
Chemotaxis assays for AWA, AWC and ASH/ADL neurons and DAF-16::mNG imaging. **(A** and **B)** Schematic of chemotaxis assays for AWA, AWC (**A**) and ASH/ADL (**B**) neurons in day 6 animals in response to 1% 2-methypyrazine, 0.5% benzaldehyde and octanol, respectively. (**A**) In AWA/AWC chemotaxis assays, day 6 animals that responded to odorants (2-methypyrazine or benzaldehyde) and migrated into region ‘A’ were grouped as AWA(+) or AWC(+). Animals that didn’t respond to odorants and migrated into ‘C’ and ‘D’ regions were grouped as AWA(-) or AWC(-). (**B**) In ASH/ADL chemotaxis assays, day 6 animals that avoided octanol and migrated into ‘E’ and ‘F’ regions were grouped as ASH/ADL(+). Animals that didn’t respond to octanol and migrated into ‘A’, ‘B’ and ‘C’ regions were grouped as ASH/ADL(-) **(C** and **D)** Function correlation analyses between AWA/AWC and ASH/ADL neurons. Data are shown as mean ± SD. Student’s *t*-test, ns, no significant difference. **(E)** Confocal images show the expression of DAF-16::mNG in ASH and ADL neurons in *daf-16(ot853)* animals. P*gpa-11*::mCherry was used as ASH/ADL marker. Scale bar: 10μm. **(F)** Confocal images show no expression of DAF-16::mNG in AMSh glia in day 1 and day 6 *daf-16(ot853)* animals. *yadIs222 [Pf53f4.13::mCherry]* was used as AMsh glia marker. Scale bar: 10μm.

**Figure S2.**
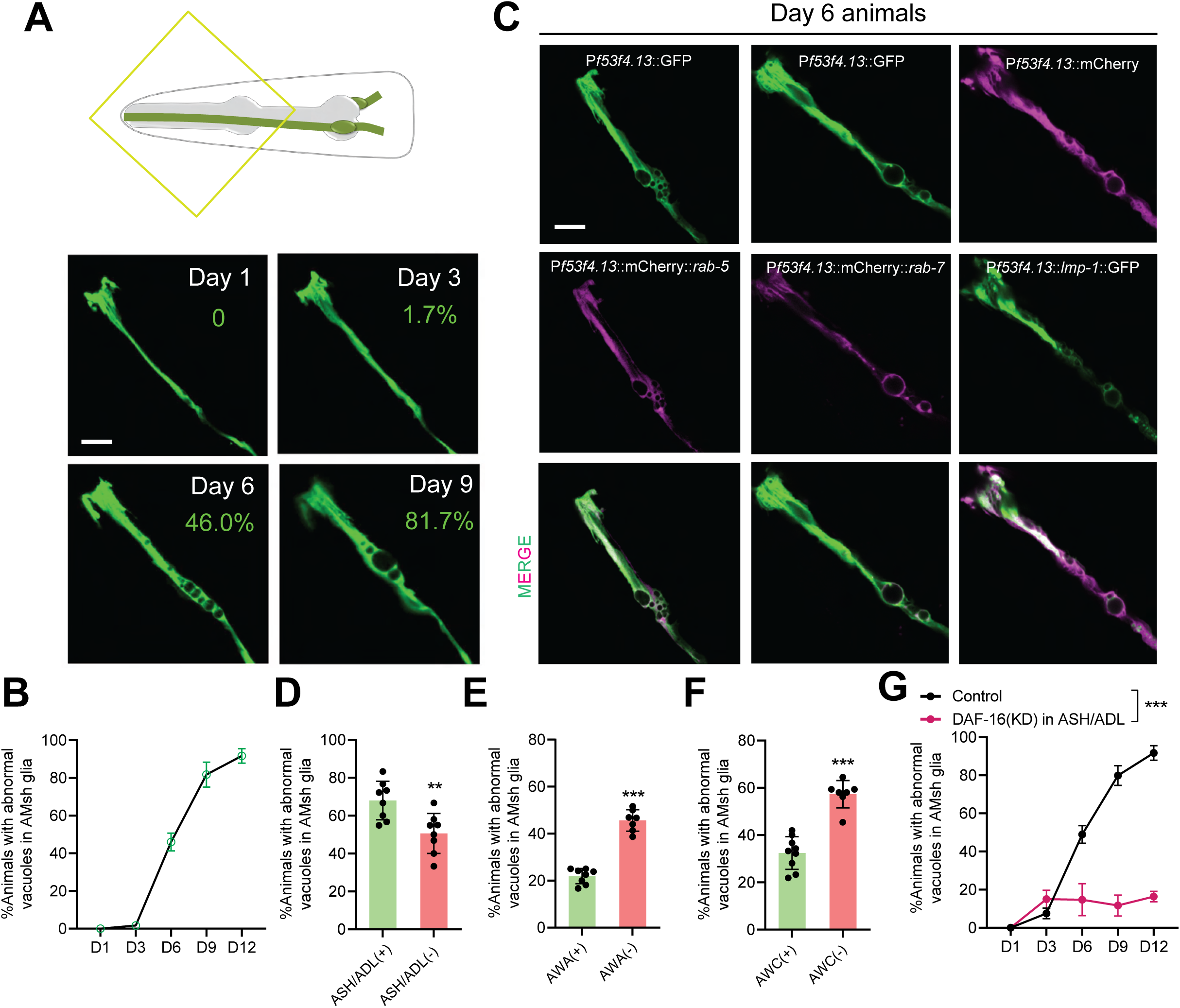
Aging-associated features in AMsh glia is regulated by ASH/ADL aging. **(A)** Abnormal vacuoles were observed in AMsh glia during aging. *yadIs48* [P*f53f4.13*::GFP] was used to imaging the morphology of AMsh glia. Scale bar: 10μm. **(B)** Quantifications of percentage of animals with abnormal vacuoles in AMsh glia during aging. **(C)** Confocal images of abnormal vacuoles in AMsh glia with early-endosome marker mCherry::RAB-5, late-endosome marker mCherry::RAB-7, and lysosome marker LMP-1::GFP. Scale bar: 10μm. **(D** to **F)** Correlation analyses between the functions of ASH/ADL (**D**), AWA (**E**), and AWC (**F**) neurons with AMsh glial aging by using day 6 animals. Data are shown as mean ± SD. Student’s *t*-test, ** P<0.01, *** P<0.001. **(G)** Quantifications of the ratio of abnormal vacuoles in AMsh glia during aging in animals with (DAF-16 (KD)) or without (Control) DAF-16 depletion in ASH/ADL neurons. Data are shown as mean ± SD. Two-way ANOVA, *** P<0.001.

**Figure S3.**
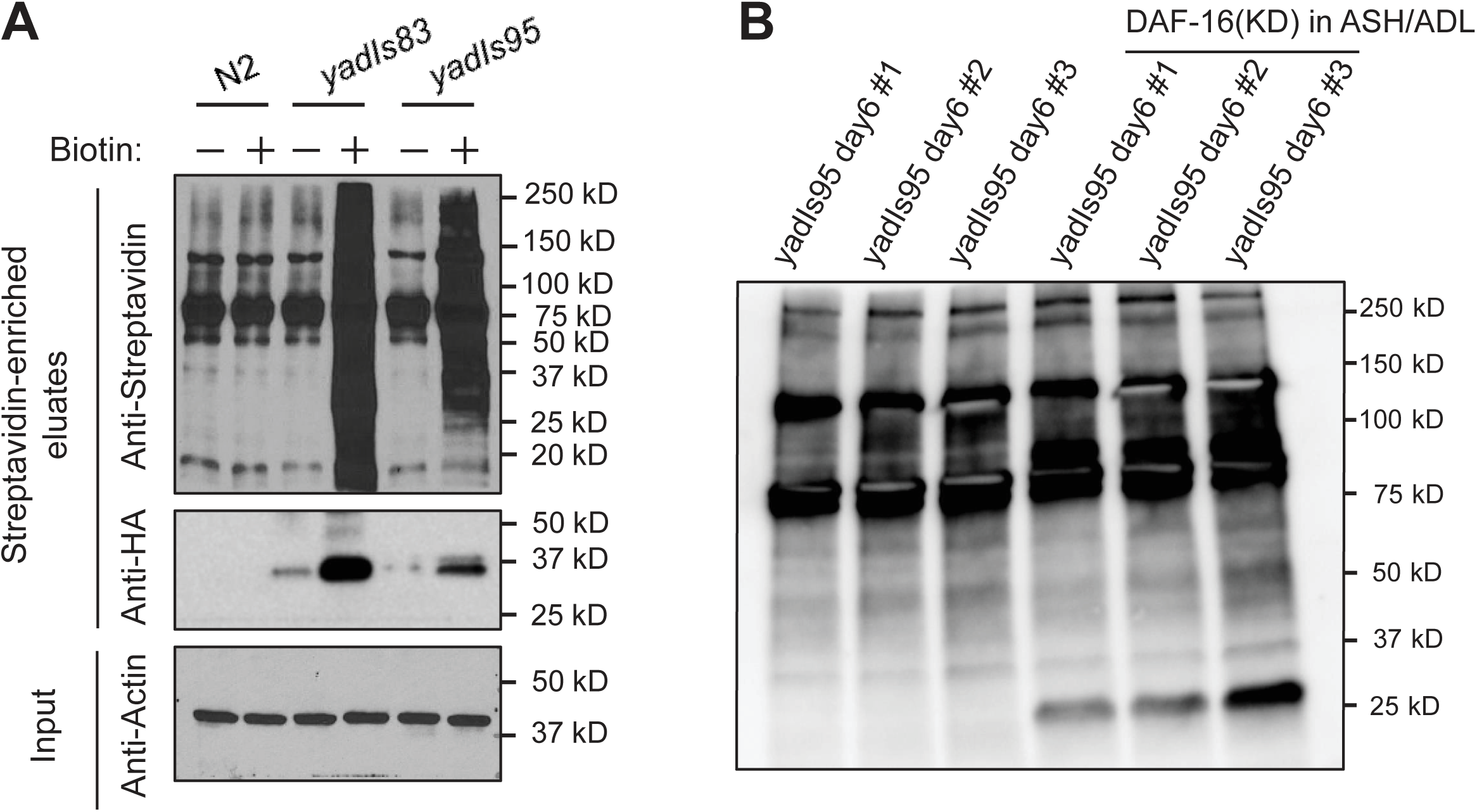
Enrichment of biotinylated proteins from AMsh glia after proximity labeling. **(A)** Enrichment of AMsh glial proteins by streptavidin magnetic beads. Wild-type (*N2*) and *yadIs83* [P*rgef-1(neuronal)*::3xHA::TurboID] animals were used as negative and positive control for proximity labeling experiments, respectively. *yadIs95* [P*f53f4.13*::3xHA::TurboID] animals express free TurboID exclusively in AMsh glia. All strains were treated with or without 100μM exogenous biotin and cultured at 25°C for 8 hours before sample collection. 50μL streptavidin-coated beads were then used for enrichment of biotinylated proteins. The protein samples were analyzed by western blotting. **(B)** Immunoblotting of enriched biotinylated AMsh glial proteins with streptavidin-HRP in transgene *yadIs95* [P*f53f4.13*::3xHA::TurboID] with or without DAF-16 depletion in ASH/ADL.

**Figure S4.**
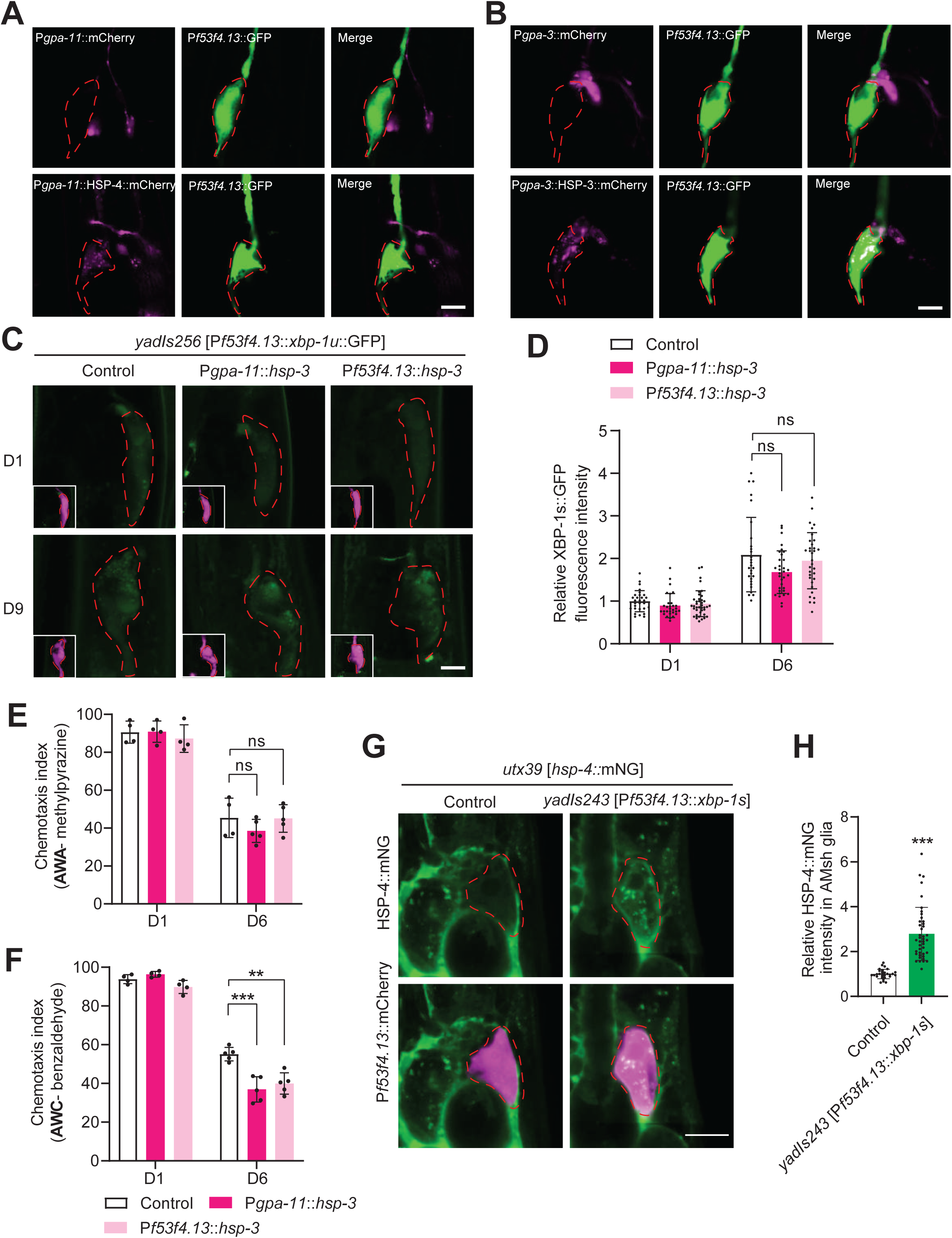
HSP-4::mCherry is transmitted from ASH/ADL neurons to AMsh glia. **(A)** Confocal images show that ASH/ADL-expressed HSP-4::mCherry but not mCherry were found in AMsh glia. HSP-4::mCherry or mCherry were specifically expressed in ASH/ADL neurons by the *gpa-11* promoter. *yadIs48* [P*f53f4.13*::GFP] was used as AMsh glia marker. Scale bar: 10μm. **(B)** Confocal images show the expression of HSP-3::mCherry or mCherry when they were specifically expressed in AMsh-channel neurons. *gpa-3* promoter was used to drive cell-specific expression in AMsh-channel neurons. *yadIs48[Pf53f4.13::GFP]* was used as a marker to label AMsh glia. Scale bar: 10 μm. **(C** and **D)** Confocal images (**C**) and quantifications (**D**) of XBP-1s::GFP intensity in control and transgenes expressing *hsp-3* in ASH/ADL neurons or AMsh glia. *gpa-11* promoter was used to drive cell-specific expression in ASH/ADL neurons. *f53f4.13* promoter was used to drive cell-specific expression in AMsh glia. *yadIs222 [Pf53f4.13::mCherry]* was used as an AMsh glia marker. Scale bar: 10μm. N>30. Data are shown as mean ± SD. One-way ANOVA, ns, no significant difference. **(E** and **F)** Results from chemotaxis assays show AWA (**E**) and AWC (**F**) neuronal functions in control and transgenes expressing *hsp-3* in ASH/ADL neurons or AMsh glia. *gpa-11* promoter was used to drive cell-specific expression in ASH/ADL neurons. *f53f4.13* promoter was used to drive cell-specific expression in AMsh glia. Data are shown as mean ± SD. One-way ANOVA, ns, no significant difference, **P < 0.01, ***P < 0.001. **(G** and **H)** Confocal images (**G**) and quantifications (**H**) of *utx39 [hsp-4::mNG::3xFlag]* show that HSP-4::mNG expression in AMsh glia is increased in animals expressing constitutively activated *xbp-1s* in AMsh glia. Scale bar: 10μm. Data are shown as mean ± SD. Student’s *t*-test, *** P<0.001.

**Figure S5.**
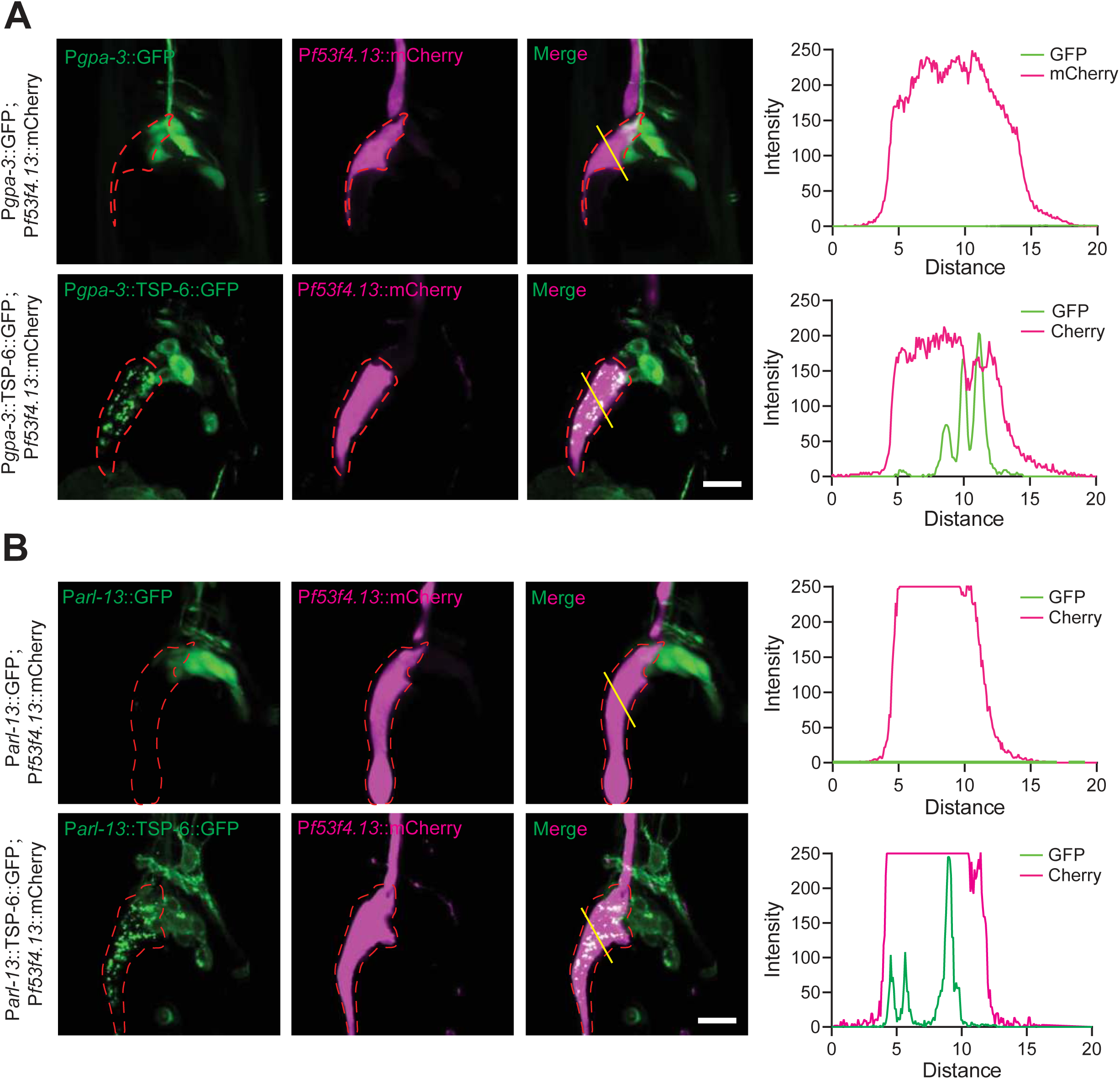
Extracellular vesicles originated from AMsh-channel neurons are transmitted to AMsh glia. **(A)** Confocal images showed that TSP-6::GFP but not GFP expressed in AMsh-channel neurons by the *gpa-3* promoter translocated into AMsh glia. The GFP and mCherry intensities along the yellow lines indicated in the figures were quantified and shown in the graphs on the right side. *yadIs222* [P*f53f4.13*::mCherry] was used as AMsh glia marker. Scale bar: 10μm. **(B)** Confocal images showed that TSP-6::GFP but not GFP expressed in amphid sensory neurons by the *arl-13* promoter translocated into AMsh glia. The GFP and mCherry intensities along the yellow lines indicated in the figures were quantified and shown in the graphs on the right side. *yadIs222* [P*f53f4.13*::mCherry] was used as AMsh glia marker. Scale bar: 10μm.

**Figure S6.**
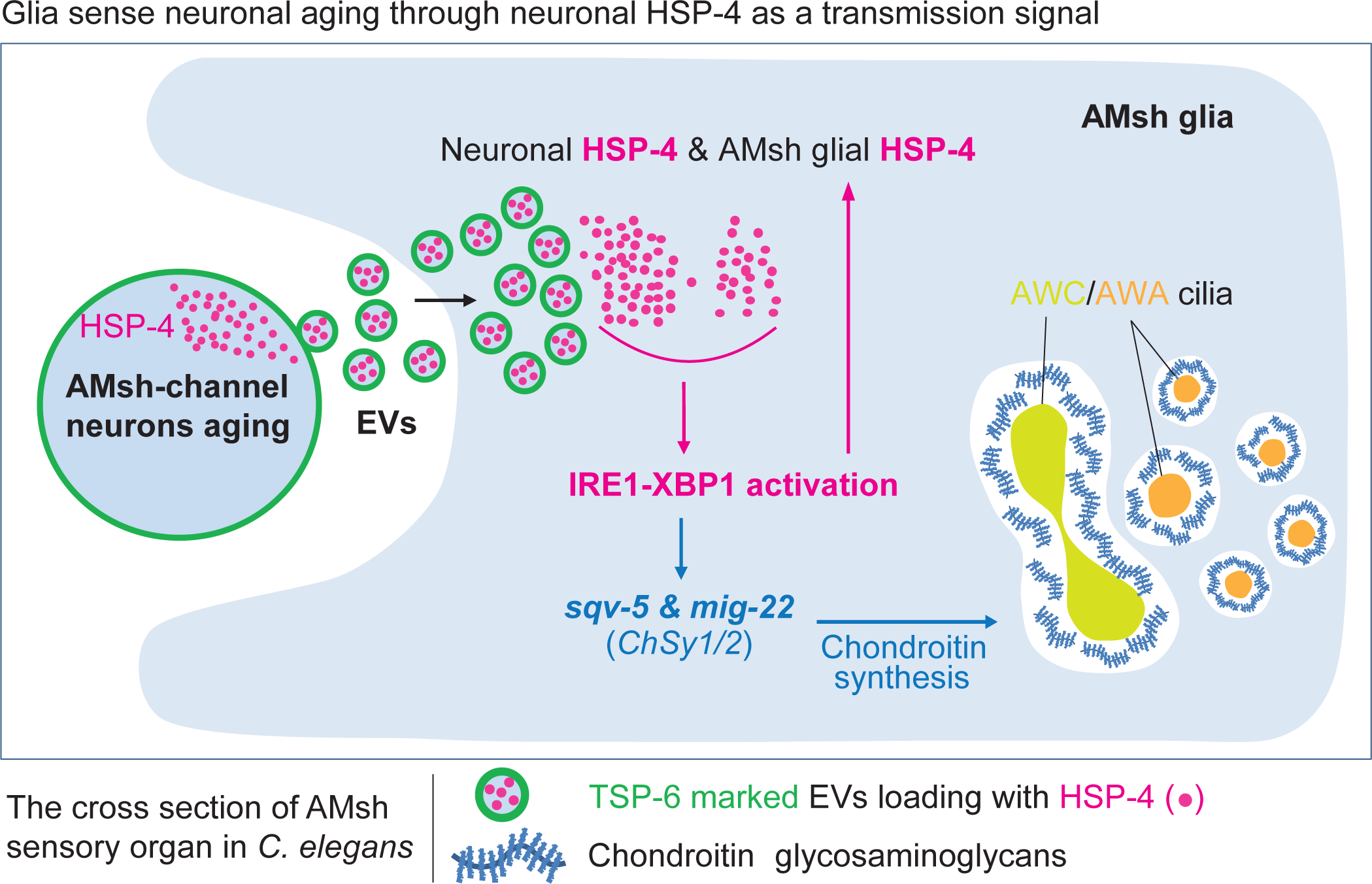
Glia sense neuronal aging through HSP-4 as a transmission signal. A model for the neuron-glia-neuron regulatory mechanism during aging, in which neuronal HSPs are transmitted to AMsh glia via EVs and activate the IRE1-XBP-1 pathway in AMsh glia to protect AMsh glia embedded neurons form aging.

**Table S1. List of strains used in this study**

**Table S2. List of plasmids used in this study**

**Table S3. Normalized data from LC-MS/MS**

## Notes

### Competing Interest Statement

The authors have declared no competing interest.

